# A promiscuity locus confers *Lotus burttii* nodulation with rhizobia from five different genera

**DOI:** 10.1101/2021.08.26.457880

**Authors:** Mohammad Zarrabian, Jesús Montiel, Niels Sandal, Haojie Jin, Yen-Yu Lin, Verena Klingl, Macarena Marín, Euan James, Martin Parniske, Jens Stougaard, Stig U. Andersen

## Abstract

Legumes acquire access to atmospheric nitrogen through nitrogen fixation by rhizobia in root nodules. Rhizobia are soil dwelling organisms and there is a tremendous diversity of rhizobial species in different habitats. From the legume perspective, host range is a compromise between the ability to colonize new habitats, where the preferred symbiotic partner may be absent, and guarding against infection by suboptimal nitrogen fixers. Here, we investigate natural variation in rhizobial host range across *Lotus* species. We find that *Lotus burttii* is considerably more promiscuous than *Lotus* japonicus, represented by the Gifu accession, in its interactions with rhizobia. This promiscuity allows *Lotus burttii* to form nodules with *Mesorhizobium, Rhizobium, Sinorhizobium, Bradyrhizobium*, and *Allorhizobium* species that represent five distinct genera. Using recombinant inbred lines, we have mapped the Gifu/*burttii* promiscuity QTL to the same genetic locus regardless of rhizobial genus, suggesting a general genetic mechanism for host-range expansion. The Gifu/*burttii* QTL now provides an opportunity for genetic and mechanistic understanding of promiscuous legume-rhizobia interactions.

## Introduction

Symbiosis with nitrogen-fixing rhizobia in root nodules enable legumes to access atmospheric nitrogen. In most cases, rhizobial entry into root cells requires recognition of rhizobial Nodulation factors (NF) signalling molecules that are secreted in response to perception of plant flavonoids (Oldroyd 2013). In turn, host membrane-bound Nod factor receptors (NFRs) initiate downstream signal transduction pathways initiating rhizobial infection and nodule organogenesis (Madsen et al. 2003; Radutoiu et al. 2003; Murakami et al. 2018). Plants produce complex mixtures of flavonoids (Liu and Murray 2016). Likewise, rhizobia secrete many different Nod factor species (D’Haeze and Holsters 2002), and both flavonoid and Nod factor pools may change dynamically over time within the same plant accession or rhizobia strain (Liu and Murray 2016; Kelly et al. 2018). This provides an intricate system that, along with bacterial effectors and exopolysaccharides and the corresponding plant detection systems, allows fine tuning of legume-rhizobium compatibility (Yang et al. 2010; Kawaharada et al. 2015; Kusakabe et al. 2020). The successful establishment of nitrogen fixation requires full compatibility of symbiotic partners. Symbiotic compatibility can affect early stages of infection, determining whether or not nodules are formed, or later stages of nodule development, affecting nitrogen-fixing efficiency or nodule senescence (Perret et al. 2000; Wang et al. 2012; Yang et al. 2017).

Intercellular infection is thought to represent a more ancient and less advanced infection mode than intracellular infection through root hairs (Sprent 2007). Specific legumes are typically infected either intra- or intercellularly. However, at least some legumes maintain genetic programs for both types of infection. These include *Sesbania rostrata*, where the infection mode can change in response to flooding (Herder et al. 2006), and *Lotus* species, where intercellular infection appears to serve as a backup function to the preferred intracellular infection route through root hair infection threads. This phenomenon was observed in *Lotus japonicus* Gifu as rare infection events of spontaneous nodules in a NF receptor deficient genetic background (Madsen et al. 2010) and has since been found in the *L. japonicus* Gifu interaction with IRBG74 (Montiel et al. 2020) and in *Lotus burttii* interactions with *Sinorhizobium fredii* HH103 and *Rhizobium leguminosarum* Norway (Acosta-Jurado et al. 2016b; Liang et al. 2019). Generally, intracellular root hair infection appears to offer more stringent scrutiny of the rhizobial partner, whereas the compatibility requirements for intercellular crack entry appears to be more relaxed (Sprent 2007; Madsen et al. 2010).

In *Lotus*, both inter- and intracellular infection depend on NF signalling (Acosta-Jurado et al. 2016b; Montiel et al. 2020), except in rare cases, if organogenesis is activated in the absence of Nod factor signalling (Madsen et al. 2010). Candidate gene approaches relying on interspecific variation in Nod factor receptors has been used to demonstrate their roles in determining compatibility (Radutoiu et al. 2007). Effectors and secretion system components that deliver effectors into plant cells also affect compatibility. For instance, the bradyrhizobial NopP effector is recognised by soybeans carrying the *Rj2* NLR resistance gene, leading to termination of infection (Sugawara et al. 2018), and the *Bradyrhizobium elkanii* NopF effector prevents infection in *Lotus japonicus*, but not in *L. burttii* (Kusakabe et al. 2020). Rhizobial effectors can also promote symbiotic interactions as exemplified by the *B. elkanii* effector Bel2-5, which confers the ability to nodulate soybeans deficient in Nod factor perception (Ratu et al. 2021). Rhizobial genes that affect Nod factor and exopolysaccharide production can also influence host range as demonstrated by the *Sinorhizobium fredii* HH1103 *mucR1, syrM, nolR* and *nodD2* mutants (Acosta-Jurado et al. 2016a). Remarkably, *syrM, nolR* and *nodD2* mutations induce a shift from inter- to intracellular infection in *L. burttii* and extend the host range to include *L. japonicus* Gifu (Acosta-Jurado et al. 2019, 2020). In addition, *Lotus* intraspecific variation was exploited to identify *Pxy* as a regulator of symbiotic compatibility downstream of exopolysaccharide signalling (Kawaharada et al. 2021). Also in *Lotus, S. fredii* HH103is able to form functional nodules on *L. burttii*, whereas it induced ineffective nodules on *L. japonicus* Gifu, and a QTL was mapped to chromosome 1 near the *Nfr1* gene (Sandal et al. 2012). Likewise, *L. burttii* was also more permissive than *L. japonicus* Gifu in the interaction with *R. leguminosarum* Norway (Grossmann et al. 2012).

Here, we investigate natural variation in rhizobial host range between the two *Lotus* species *L. japonicus* Gifu and *L. burttii*, focusing on host control of symbiotic compatibility.

## Materials and Methods

### Nodulation phenotyping

For germination, seeds were scarified and surface sterilized with 0.5% sodium hypochlorite for 15 minutes and rinsed several times with distilled water. Seeds were kept in sterile water for 1 hour before sowing on wet filter paper. The germinated plants were transferred and grown on 1/4 Broughton and Dilworth (B&D) medium (Broughton and Dilworth 1971) where the surface of the agar slope was covered with filter paper. Rhizobia were grown in yest agar medium (YAM) except for *R. leguminosarum* strains that were grown in tryptone yeast medium (TY). The strains were diluted to an OD600 of 0.02 before inoculating with 50 µl per plant by pipetting the suspension directly on the root. The nodulation phenotype was recorded at 35 days post-inoculation.

### QTL Analysis

We used recombinant *L. japonicus* Gifu x *L. burttii* recombinant inbred lines (RILs) (Sandal et al. 2012; Shah et al. 2016). For rough mapping, 18 RILs with balanced genotypes were chosen (**Supplemental dataset 1**). Genotype and phenotype data were imported into R/qtl version 3.4.2 (R Project for Statistical Computing, www.r-project.org/) using the read.cross command. After converting to RIL format, the genetic map and missing genotype values were estimated using est.map and mqmaugment, respectively. Multiple QTL Mapping (MQM) was then conducted using 1000 permutations to determine significance thresholds.

### Hairy root transformation with *Nfr1* constructs

The *L. burttii Nfr1* (BinG1) construct was based on the *L. japonicus Nfr1* complementation construct, (carrying the entire *LjNfr1* gene driven by its own promoter), which was modified using standard cloning techniques and transferred into the pIV10 integration vector (AM235368). The construct was integrated into the *Agrobacterium rhizogenes* strain AR12 (Hansen et al. 1989). BinG1 was constructed as follows: a DNA fragment from position 4090 to position 4993 of the *Nfr1* gene (AJ575246/AJ575247) was substituted by the corresponding fragment from *L. burttii* produced by PCR. The two *L. japonicus* Gifu/*L. burttii* polymorphisms identified in the *Nfr1* extracellular region are both contained within the *L. burttii* fragment included in the *BinG1* construct.

### Genotyping and phenotyping of F1 progeny of *L. burttii* and *L. japonicus* crosses

Crosses using *L. burttii* as mother and *L. japonicus* Gifu as father, or the converse, were generated. Seeds *of L. burttii* B-303, *L. japonicus* Gifu B-129 and the F1 progeny were scarified and surface-sterilized as described earlier. Six-day old seedlings were transferred into tulip shaped Weck jars (Weck 745) containing 300 ml sterilized sand-vermiculite mixture supplemented with 40 ml of FAB medium. After two days, each plant was inoculated with 1 ml of a *R. leguminosarum* Norway suspension (OD_600_ = 0.005) or 1 ml FAB medium as a mock control. Plants were grown under a long-day photoperiod for six weeks and phenotyped using a MZ16 FA stereomicroscope (Leica). For genotyping, genomic DNA was extracted from leaves lysed in liquid nitrogen. Lysates were suspended in 500 μl extraction buffer (2% w/v CTAB, 1.42 M NaCl, 20 mM EDTA, 100 mM Tris-HCl, pH8) supplemented with 3.1 μl beta-mercaptoethanol and incubated at 65 ° C for 20 min. Suspensions were mixed with 300 μl chloroform and centrifuged at 14,000 rpm for 5 min. Supernatants were mixed with 1/10 volume of 3 M NaOAc and centrifuged at 14,000 rpm for 15 min. Pellets were washed two times with 70% ethanol, air-dried and re-suspend in 50 μl distilled water. The DNA was used as template in PCR reactions with the *Lotus* marker TM1203 (forward: TTGAATAAGGCTCATAGATCC, reverse: CTTCAGTTTGGGTTTCAAGC) (Sato et al., 2001) and verified by agarose gel electrophoresis.

## Results

### *L. burttii* nodulates with rhizobia from five different genera

It was previously reported that *S. fredii* HH103 forms functional nodules on *L. burttii* but not on *L. japonicus* Gifu (Sandal et al. 2012). In order to determine if *L. burttii* is generally more permissive in its symbiotic interactions than Gifu, we examined the nodulation phenotypes of Gifu and *L. burttii* with a wide range of rhizobia from different genera including *Sinorhizobium, Azorhizobium, Bradyrhizobium, Rhizobium, Allorhizobium* and *Mesorhizobium* (**Figure 1**). These distantly related rhizobial species produce NF with different chemical modifications at the nonreducing and reducing ends (D’Haeze and Holsters 2002; Beck et al. 2010; Renier et al. 2011) (**Table 1;**). We observed large variation in nodule numbers and structures, which included nodule primordia “bumps”, white nodules and in some cases small or more developed pink nodules, indicative of nitrogen fixation (**Figure 2 A-D**). Among the 42 strains tested, only the cognate *Lotus* symbiont *M. loti* R7A induced development of pink nodules in both Gifu and *L. burttii* (**Figure 2A and D**). At the other extreme, *Sinorhizobium meliloti* nodulated neither. *Sinorhizobium fredii* NGR234, which is compatible with a very broad range of legumes (Pueppke 1999), was unique in promoting a larger number of pink nodules in Gifu than in *L. burttii* (**Figure 2D; Supplementary Figure S1**). None of the remaining 39 strains formed pink nodules with Gifu. In contrast, 30 out of the 39 strains formed at least some pink nodules with *L. burttii* (**Figure 2A and D**). The 30 strains that nodulate *L. burttii* comprise five of the genera tested, indicating that the host range of *L. burttii* is broad and that *L. burttii* is considerably more promiscuous in its symbiotic interactions than Gifu, regardless the diverse composition of the NF produced by the rhizobial strains (**Table 1**).

**Table 1.** NF structure in rhizobial strains from different genera. Composition and chemical decorations in the nonreducing and reducing end of NF produced by rhizobial species used in this study (**\***) and closely related strains. Table adapted from D’Haeze and Holsters (2002). Abbreviations: Ac, acetyl; Ara, arabinosyl; Cb, carbamoyl; Fuc, fucosyl; Me, methyl; S, sulfate ester; H, hydroxide. The colour code reflects the phylogenetic proximity of the species shown in Figure 1.

| Rhizobial strain | Nod factor substitutions |  |  |
| --- | --- | --- | --- |
|  | Acyl group | Non-reducing end | Reducing end |
| ***Azorhizobium* Sp.** |  |  |  |
| * *A. caulinodans* ORS571 | C18:1,C18:0 | Me, Cb, H | Fuc, Ara, Me, H |
| ***Sinorhizobium* Sp.** |  |  |  |
| * *Sinorhizobium* NGR234 | C18:1,C16:0 | Me, Cb, H | 3-O-S-2-O-MeFuc , Me, H |
| * *S. fredii* HH103 | C16:0,C16:1 | H | 2-O-MeFuc, Fuc, Me, H |
| * *S. fredii* USDA257 | C18:1 | H | 2-O-MeFuc, Fuc, Me, H |
| *S. teranga* bv. *acaciae* ORS1073 | C16:0,C18:0,C18:1 | Me, Cb, H | S, H |
| *S. teranga* bv. *sesbaniae* | C16:0,C18:1 | Me, Cb, H | Fucy, Ara, Me, H |
| * *S. meliloti* RCR2011 | C16:1,C16:2,C16:3 | Ac, H | Me, H |
| ***Rhizobium* Sp.** |  |  |  |
| *R. leguminosarum* bv. *viciae* A1 | C16:0,C16:1,C18:0 | Ac, H | Ac, Me, H |
| *R. leguminosarum* bv. *viciae* RBL5560 | C18:4,C18:1,C18:0 | Ac, H | Me, H |
| *R. leguminosarum* bv. *viciae* TOM | C18:4,C18:1 | Ac, H | Ac, Me, H |
| ***Mesorhizobium* Sp.** |  |  |  |
| * *M. loti* R7A | C16:0,C16:1,C18:0 | Me,Cb, Fuc, H | Ac, Fuc, H |
| ***Bradyrhizobium* Sp.** |  |  |  |
| *Bradyrhizobium* strains | C18:1,C18:2 | Me, Cb, H | 3-O-S-2-O-MeFuc, H |
| * *Bradyrhizobium* ORS285 | C18:1 | Me, Cb, H | 2-O-MeFuc, H |

**Figure 1.**
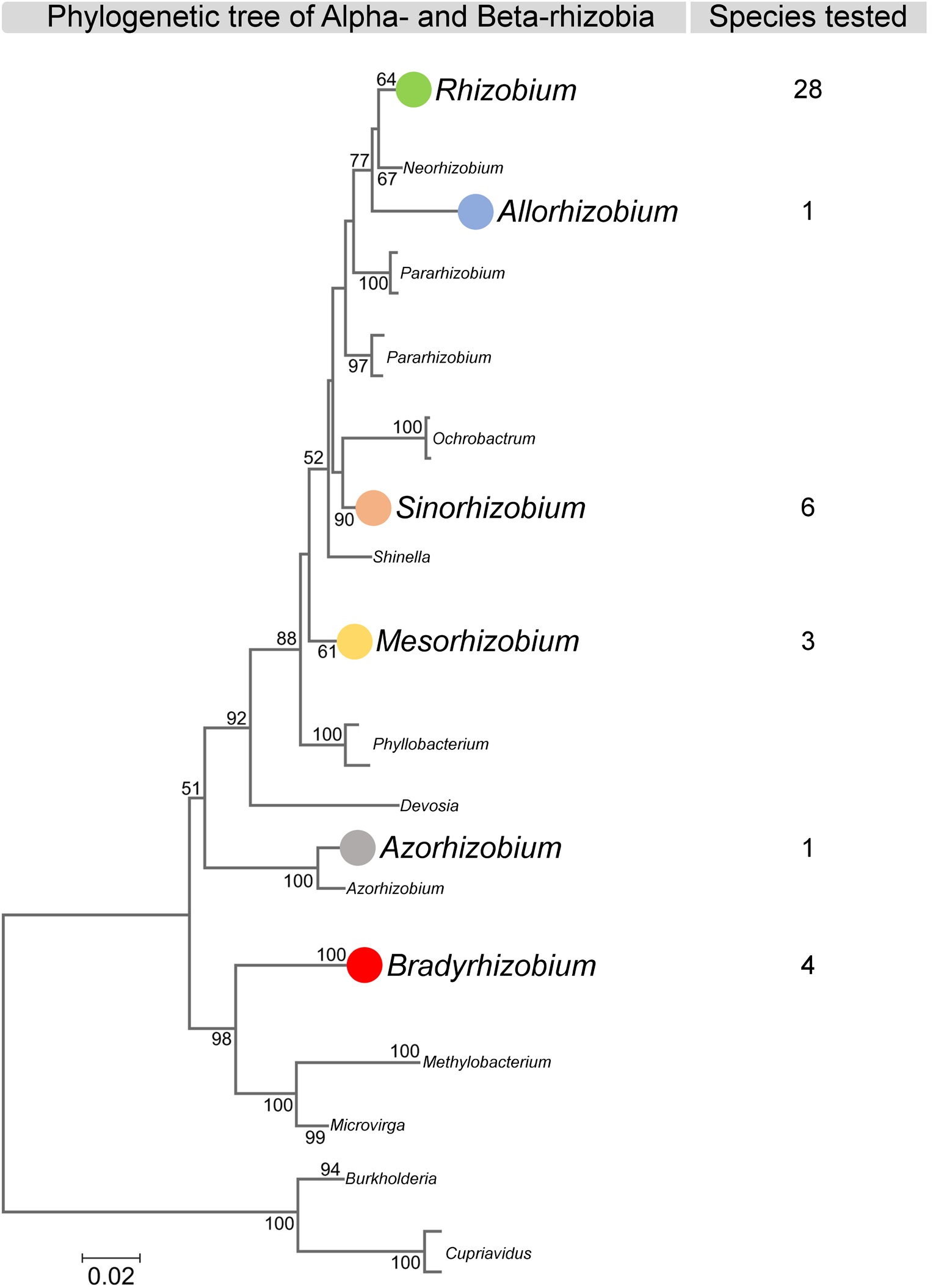
Phylogenetic distribution of rhizobia used in this study. Phylogenetic tree of Alpha and Beta-rhizobia adapted from Sprent et al. (2017) to highlight the number of species used in this work from each rhizobial genus to evaluate the nodulation capacity of *L. japonicus* Gifu and *Lotus burttii*.

**Figure 2.**
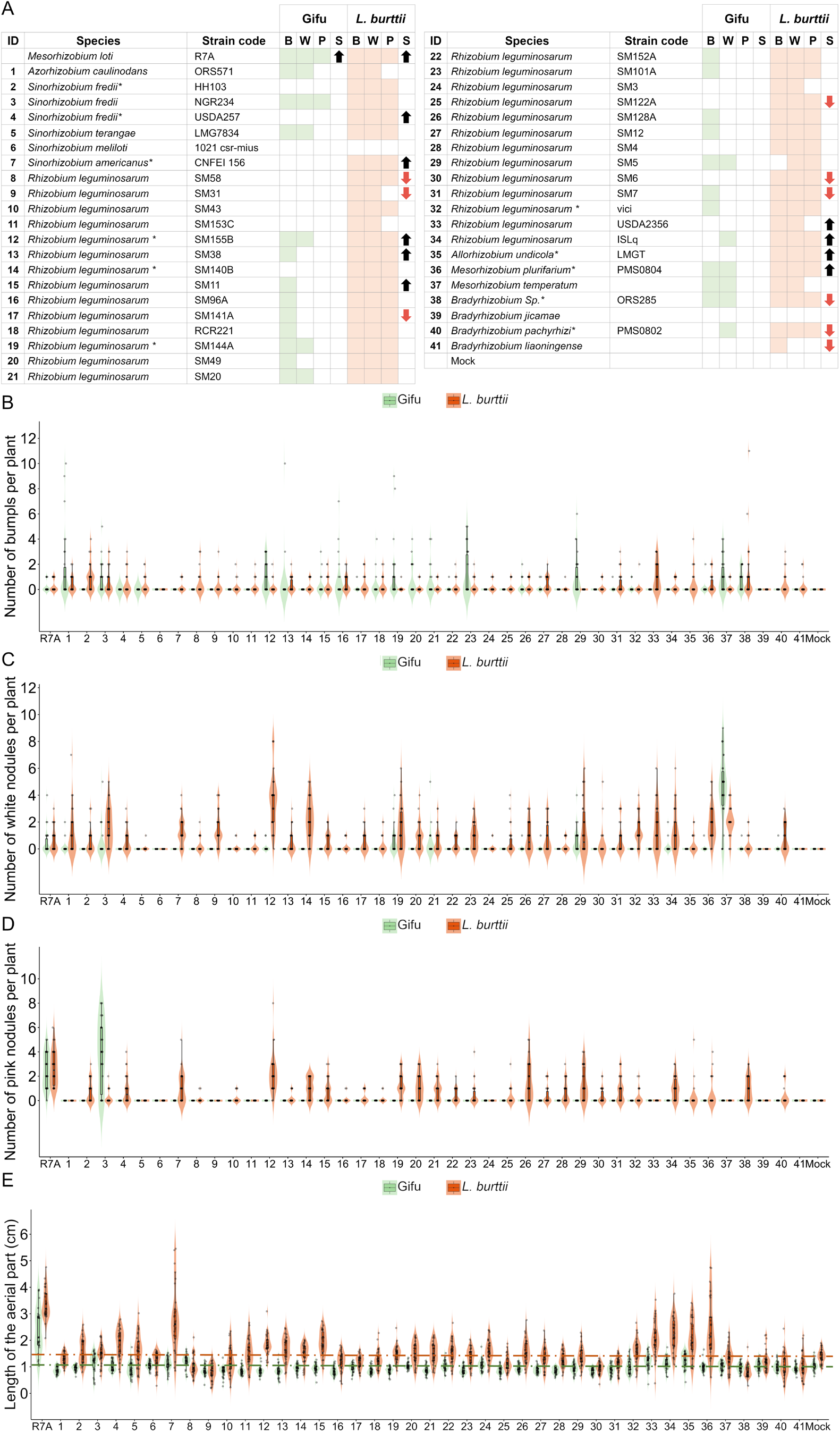
Nodulation and shoot phenotypes. **A**, Presence (filled rectangles) or absence (unfilled rectangles) of bumps (B), white nodules (W), and pink nodules (P) in Gifu and *L. burttii* plants at 5 wpi with 42 different rhizobial strains. In column S, the black and red arrows indicate a significant increase or decrease of the shoot length respect to mock-treated plants, respectively. Student’s t test; P < 0.01. **B-E**, Violin dot plots showing the number of bumps, white nodules, pink nodules and shoot length in Gifu and *L. burttii* harvested at 5 wpi with the rhizobial indicated in 2A. Dashed lines in green and orange highlight the average shoot length in mock-treated plants of Gifu and *L. burttii*, respectively.

### *L. burttii* allows nodulation with inefficient nitrogen fixers

Efficient nitrogen fixation by rhizobia in fully developed nodules has a positive impact on the plant host, reflected in a more vigorous growth of the aerial part (Lindstrom and Mousavi, 2019). Accordingly, both Gifu and *L. burttii* plants nodulated by *M. loti* R7A showed a significantly greater shoot length compared to mock-treated plants (**Figure 2A and E**). None of the 41 remaining Gifu-rhizobial associations had a positive impact on shoot length. In contrast, *L. burttii* growth was significantly enhanced by 10 different rhizobial species, nine of which developed pink nodules. However, *R. leguminosarum* strain USDA2356 (strain #33), which did not induce any pink nodules, resulted in significantly longer shoot lengths than the mock control (**Figure 2A and E**), suggesting a growth promoting effect independent of nitrogen fixation.

Despite formation of pink nodules, seventeen *L. burttii*-rhizobia interactions resulted in plants displaying similar shoot length to mock-treated plants, whereas a significant reduction of plant growth was observed in plants nodulated by nine rhizobial species (**Figure 2A and E**). These results show that although a wide range of rhizobial species developed pink nodules with *L. burttii*, most of them did not have a positive impact on growth and instead resulted in neutral or negative outcomes. This finding prompted us to further analyse the correlation between rhizobial colonization and plant growth. We used a subset of 11 *L. burttii*-rhizobia combinations, which included strains from the five different genera that formed pink nodules with *L. burttii*. For each of the 11 different *L. burttii*-rhizobia associations, only the shoot length of plants with fully developed pink nodules was compared to mock-treated plants and to plants nodulated by *M. loti* R7A (**Figure 3A**). Additionally, the nodule structure and bacteroid occupancy in the nodule cells were visualized by light microscopy (**Figure 3B-M)**. Only *L. burttii* plants harbouring pink nodules colonized by *S. americanus* and *A. undicola* (strains #7 and #35) showed comparable shoot growth to plants nodulated by *M. loti* R7A (**Figure 3A**). In contrast, plants nodulated by *Bradyrhizobium* Sp. ORS285 (strain #38) exhibited a significant reduction in the shoot length, while the remaining *L. burttii*-rhizobia interactions did not affect the length of the aerial part (**Figure 3A**). Nodule sections revealed successful rhizobial colonization by all 11 strains tested, though to different extents. *L. burttii* nodules were heavily colonized by *R. leguminosarum* SM140B, *R. leguminosarum* SM144A and *Bradyrhizobium* ORS285 (**Figure 3G, H and L**), but none of these strains had a positive effect on plant shoot length (**Figure 3A**). These results confirm that the permissive nature of *L. burttii*, which allows successful nodulation by a broad range of rhizobial species, does not always result in growth promotion.

**Figure 3.**
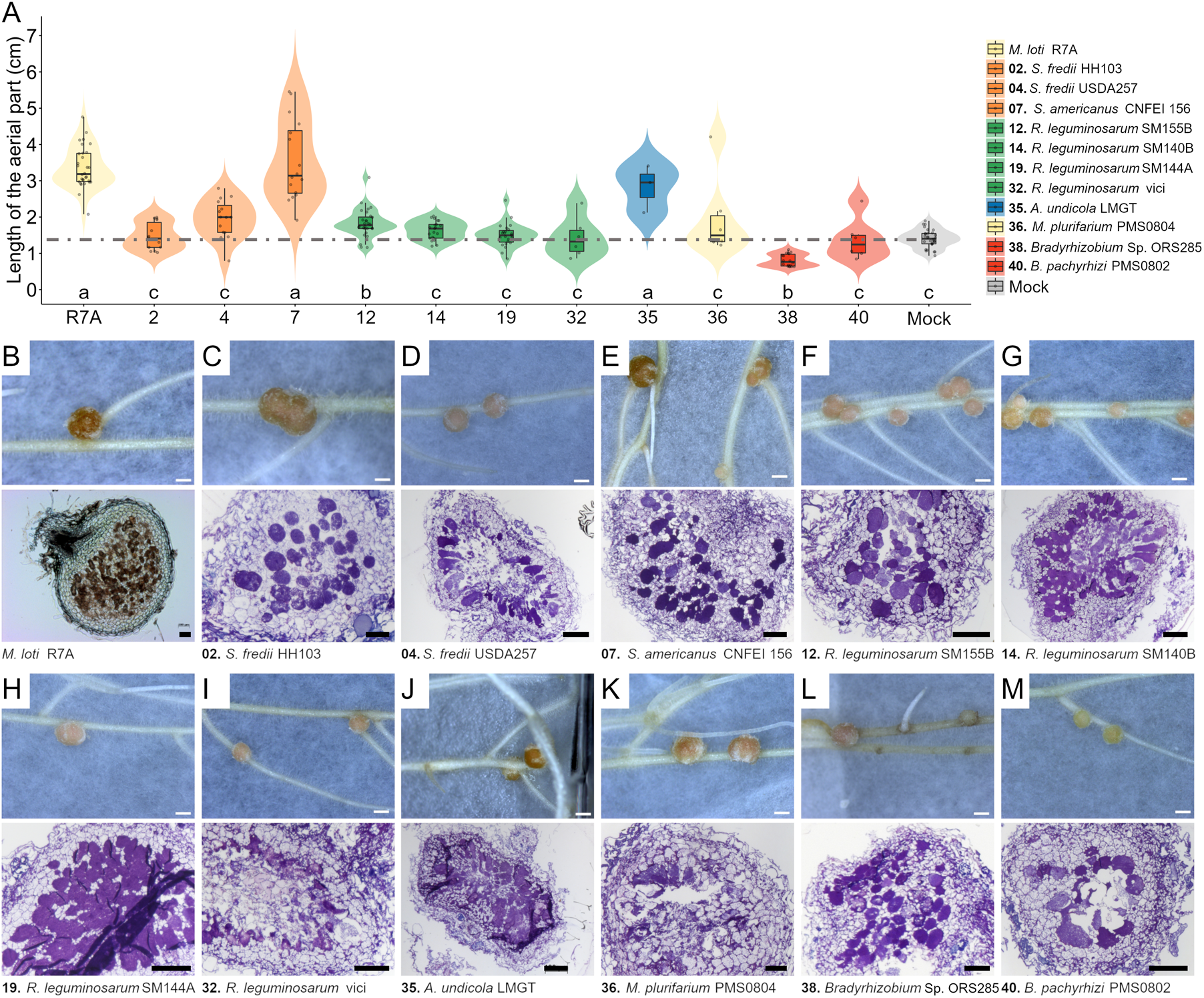
Shoot length and nodule histology of *L. burttii* plants. **A**, Violin dot plots showing the shoot length of *L. burttii* plants with pink nodules at 5 wpi with rhizobial species from different genera. Letters a and c, below the violin plots, indicate non-significant differences between the shoot length compared to mock-treated or R7A-inoculated plants, respectively; b, means significant difference to mock-treated and R7A-inoculated plants. ANOVA, P-Tukey < 0.01. The colour code reflects the phylogenetic proximity of the species shown in Figure 1. Dashed grey line highlights the average shoot length in mock-treated plants. **B-M**, Nodule histology with representative images of pink nodules developed in *L. burttii* by different rhizobial strains. Scale bar, 1 mm for nodules and 100 µm nodule sections.

### A single genetic locus near *Nfr1* confers *L. burttii* promiscuity

With the exception of *S. fredii* NGR234, Gifu only formed large, pink nodules with its cognate efficient nitrogen fixer *M. loti* that contributed positively to plant growth. The situation was much more complex for *L. burttii*, which engaged in many more different interactions with variable outcomes for the plant. In order to understand the genetics underlying *L. burttii* promiscuity, we inoculated 18 Gifu x *L. burttii* RILs (Sandal et al., 2012) with the 11 diverse strains mentioned above and *R. leguminosarum* Norway, which nodulates *L. burttii* roots intercellularly but does not form nodules with Gifu (Grossmann et al. 2012; Liang et al. 2019) (**Figure 3; Supplementary dataset 1**).

We scored the average number of pink nodules for each RIL-rhizobium combination (**Supplemental dataset 1**) and analysed the resulting data using R/qtl. TM0002, the marker previously reported to be associated with the *S. fredii* HH103 Gifu/*L. burttii* QTL (Sandal et al. 2012), was identified as the highest peak in the QTL analyses for all 12 strains (**Figure 4**). Interestingly, the single locus that appears to be responsible for allowing *L. burttii* nodulation with all 12 strains tested was located on chromosome 1, near the Nod Factor Receptor gene *Nfr1*.

**Figure 4.**
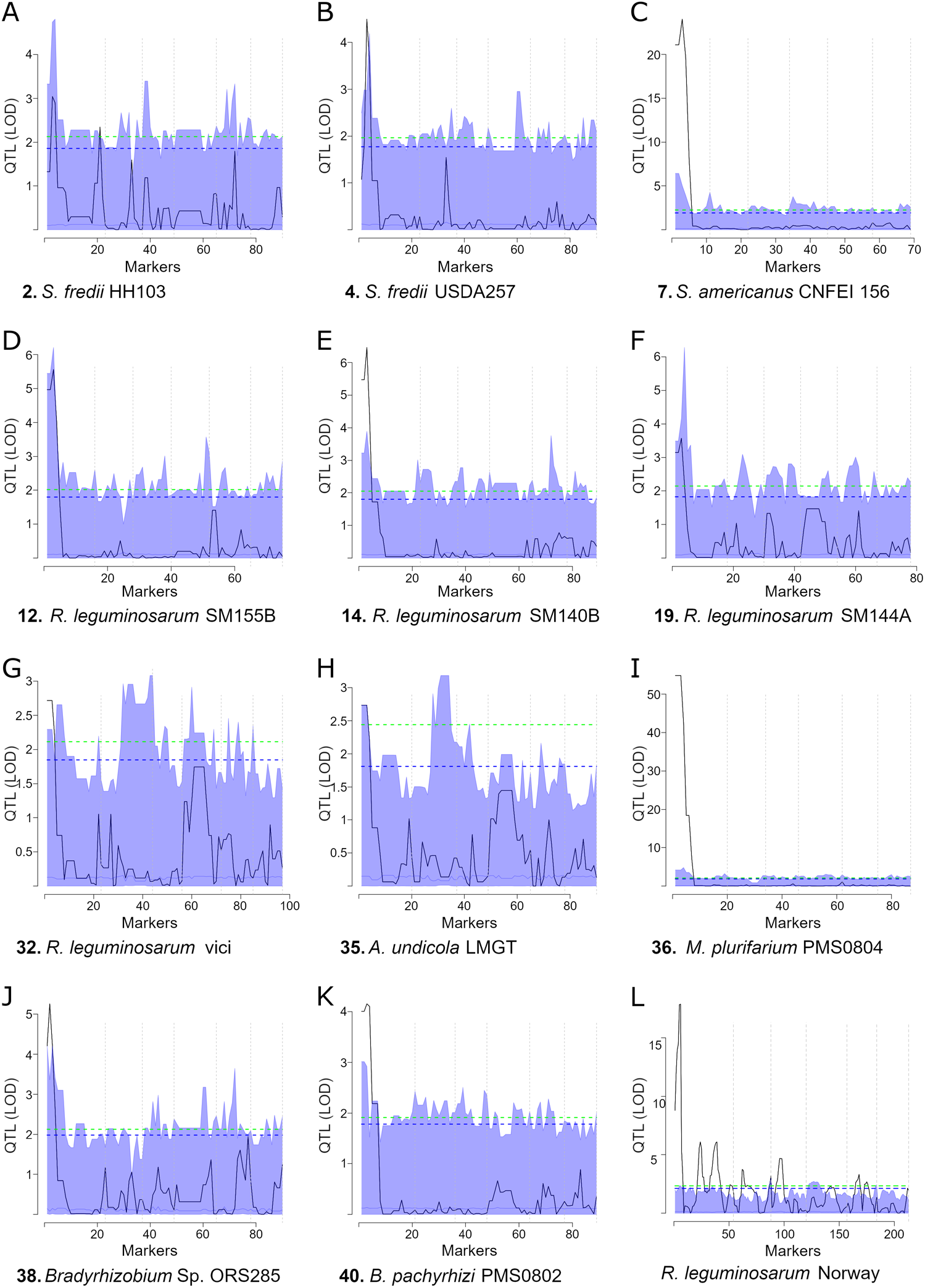
QTL analysis of Gifu x *L. burttii* RIL nodulation. The trait analysed is the average number of pink nodules. A: *S. fredii* HH103. B: *S. fredii* USDA257. C: *S. americanus* CFEI156. D: *R. leguminosarum* SM155B. E: *R. leguminosarum* SM140B. F: *R. leguminosarum* SM144A. G: *R. leguminosarum* vici. H: *A. undicola* LMGT. I: *M. plurifarium* PMS0804. J: *Bradyrhizobium* sp. ORS285. K: *B. pachyrhizi* PMS0802. L: *R. leguminosarum* Norway.

### A *L. burttii*/Gifu *Nfr1* domain swap construct complements the *nfr1* mutant in a Gifu background, but does not extend its host range

Our analysis revealed that an *Nfr1*-linked locus, or *Nfr1* itself, is involved in determining permissiveness towards diverse rhizobial species in *L. burttii*. We sequenced the extracellular region of the *L. burttii Nfr1* gene and found two mis-sense substitutions compared to the Gifu gene (T124A and Y213D, Gifu/*L. burttii*). This made *Nfr1* a strong candidate gene for explaining the capacity of *L. burttii* to establish symbiotic associations with multiple rhizobial genera. To examine this possibility, we created a BinG1 domain swap construct where the *Nfr1* extracellular domain from Gifu was replaced by the corresponding fragment from *L. burttii Nfr1*. This construct was introduced into Gifu wild type and *Ljnfr1-1* mutant backgrounds using *Agrobacterium rhizogenes* hairy root transformation. The *BinG1* construct was functional and rescued Gifu *nfr1-1* nodulation with *M. loti* R7A (**Figure 5**). In contrast, it did not enable either the Gifu wild type or the Gifu *nfr1-1* mutant to nodulate with *S. fredii* HH103 (**Figure 5**). A functional *L. burttii Nfr1* extracellular domain in the Gifu background was thus not sufficient for extending the rhizobial host range of Gifu.

**Figure 5.**
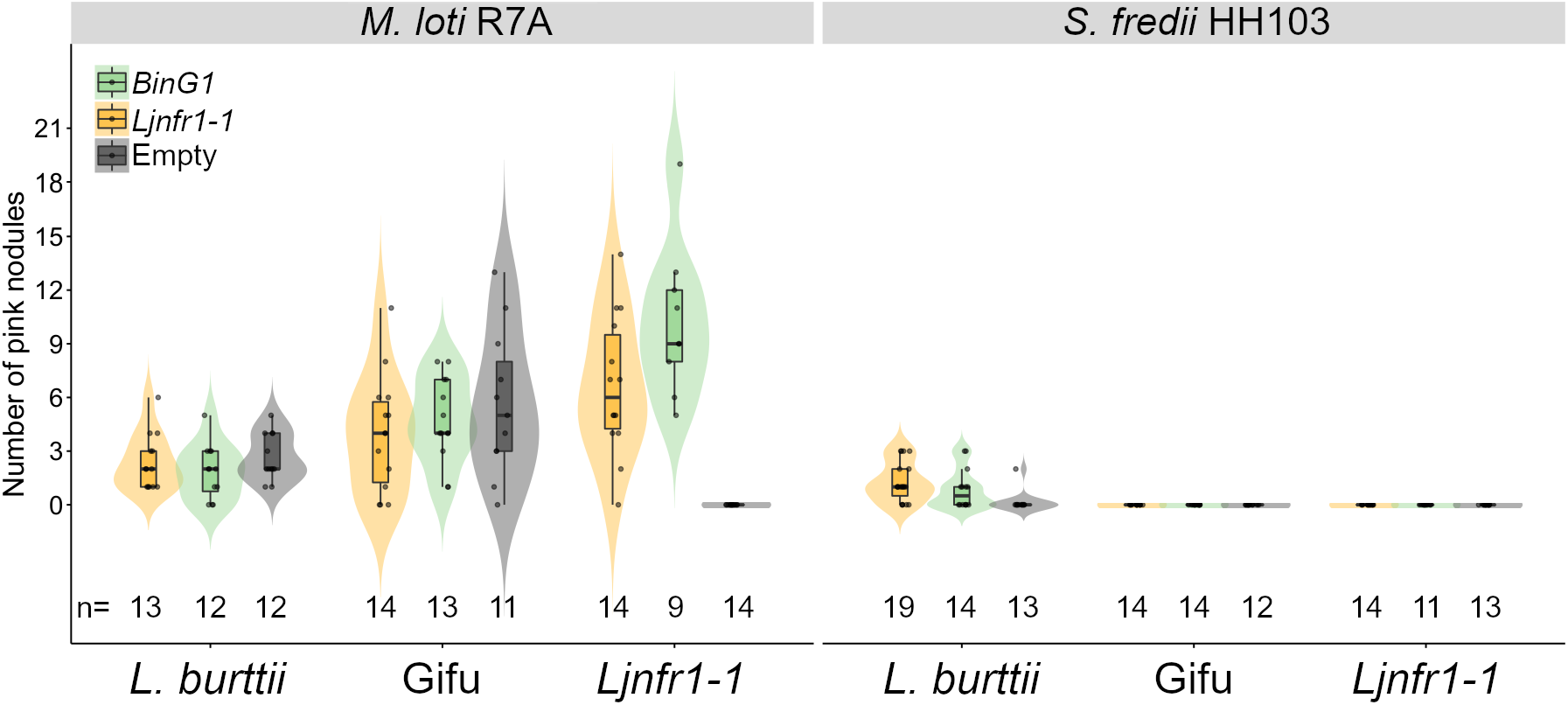
Influence of the *Nfr1* genotype on nodulation with *S. fredii* HH103. Violin dot plots show the number of pink nodules found in *L. burttii*, Gifu and the *Ljnfr1-1* mutant transformed with the *BinG1*and *LjNfr1* constructs or the empty vector at 5 wpi with *M. loti* R7A and *S. fredii* HH103. *BinG1, Nfr1* extracellular domain of Gifu replaced by the corresponding fragment from *L. burttii Nfr1* expressed under *LjNfr1* promoter. *LjNfr1*, Gifu *Nfr1* sequence expressed under its native promoter. The number of plants tested is indicated below the violin plots.

### Nodulation of *L. burttii* with *R. leguminosarum* Norway is a dominant trait

Recently it was shown that *Rhizobium leguminosarum* Norway colonizes *L. burttii* roots intercellularly (Liang et al. 2019) but is unable to infect *L. japonicus* (Grossman et al. 2012). This report along with the data presented in this study confirms the wide host range of *L. burttii* with diverse rhizobial species. To assess if this nodulation promiscuity is a recessive or dominant trait, the F1 progeny of crosses between *L. burttii* and *L. japonicus* Gifu were inoculated with *R. leguminosarum* Norway-GFP. Gifu remained without nodules six weeks post inoculation. The F1 progeny developed nodules regardless of the parental combination (**Figure 6A**) and all phenotyped F1 plants were heterozygous (**Figure 6B**). These results show that the nodulation phenotype of *L. burttii* is dominant.

**Figure 6.**
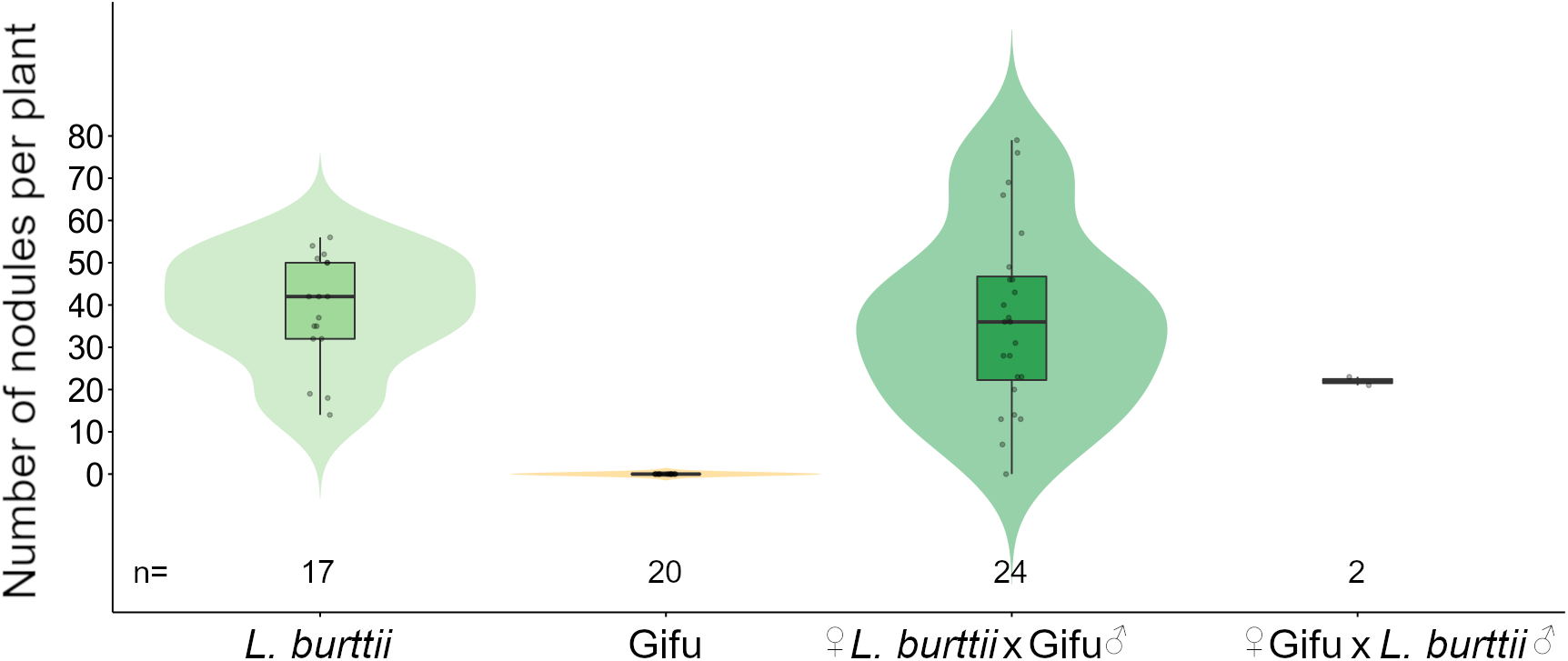
*L. burttii* x *L. japonicus* Gifu F1 genotype and nodulation phenotype with *R. leguminosarum* Norway. A, Violin dot plots show the nodule numbers at 6 wpi with *R. leguminosarum* Norway in *L. burttii, L. japonicus* Gifu and the F1 progeny from crosses between these two genotypes. The number of plants tested are described below the violin dot plots. B, Agarose gel with PCR products amplified with a set of primers for the *Lotus* power marker TM1203 using as template DNA isolated from *L. burttii* (Lb), *L. japonicus* Gifu (Lj Gifu) and the F1 progeny from their crosses (Lb x Lj Gifu).

## Discussion

The legume-rhizobia symbiosis is an illustrative example of a highly specific and stringent molecular dialogue between symbiotic partners. Despite this fact, it is known that certain rhizobial species such as *S. fredii* NGR234 is able to establish symbiotic associations with diverse legumes from distant phylogenetic genera (Pueppke and Broughton, 1999). In the last decades, both rhizobial and plant components defining the legume-rhizobia compatibility have been identified (Walker et al. 2020). This study shows that a single locus in *L. burttii* is responsible for promiscuous interactions across rhizobial genera.

### *L. burttii* a promiscuous symbiotic partner engaged in beneficial and detrimental associations

*L. burttii* is related to *L. japonicus*, a model legume with extensive genetic and transcriptomic resources (Fukai et al. 2012; Urbański et al. 2012; Mun et al. 2016; Małolepszy et al. 2016; Kamal et al. 2020). Unlike *L. japonicus*, which seems to be nodulated by a narrow range of rhizobial partners, *L. burttii* established 30 associations with diverse rhizobia that culminated in the formation of pink nodules. Excluding the *L. burttii*-*M. loti* interaction, only nine rhizobial species had a marginal but significantly positive contribution to host plant growth while the rest of the rhizobial interactions either did not affect or had a negative impact on *L. burttii* growth. Except for *R. leguminosarum* Norway (Liang et al. 2018), the rhizobial strains used in this study are efficient nitrogen fixers in the symbiotic associations with their cognate plant hosts (Cavassim et al. 2020; Moeskjær et al. 2021; Pueppke and Broughton 1999; Dreyfus and Dommergues 1981; Buendia-Claveria et al. 1989; Lajudie et al. 1994; Mora et al. 2014; Lajudie et al. 1998a, 1998b; Gao et al. 2004; Ramírez-Bahena et al. 2009; Gao et al. 2015). The suboptimal outcomes of many of these *L. burttii*-rhizobial associations may be caused by a delay in the nodulation kinetics, as occurs in the *L. burttii*-*S, fredii* HH103 symbiosis (Acosta-Jurado et al. 2016b). However, additional compatibility elements, acting at later stages of nodulation could be required for efficient nodule functioning and nitrogen fixation (Walker et al. 2020). Similar examples of inefficient legume-rhizobia symbioses have been documented in *Medicago* spp. *S. meliloti* 1021 is an efficient nitrogen-fixer in *Medicago sativa* nodules and is also able to form pink nodules with the *M. truncatula* accessions A17 and R108, but with poor nitrogen fixation performance (Terpolilli et al. 2008; Kazmierczak et al. 2017). Similarly, ineffective mutants of *S. meliloti* were comparable to their effective counterparts in colonizing *M. sativa* nodules (Amarger 1981). However, legumes possess mechanisms to reward or penalize the effectiveness of their rhizobial partners hosted within nodule cells. In soybean nodules, where inefficient nitrogen fixation was mimicked by substituting nitrogen for argon, the population and growth of rhizobia was drastically lower with respect to nodules where nitrogen fixation was performed efficiently (Kiers et al. 2003). Likewise, four-generation experiments conducted with twelve *M. truncatula* genotypes inoculated with a mixture of three rhizobial strains from their native range revealed an increase in the relative frequency of more beneficial rhizobial strains, estimated by the nodule number and size (Heath and Tiffin 2009).

A relatively low capacity of legumes to colonize new habitats seems to be related to a scarce presence of compatible symbionts (Parker 2001). This idea is supported by the high invasiveness or certain woody legumes that possess a broad compatibility with diverse rhizobial strains (Richardson et al. 2000). A comparative study among congeneric acacias revealed that those considered invasive associate with a significantly greater number of rhizobial strains than the natural and non-invasive acacias in Australia (Klock et al. 2015). However, studies conducted in distinct geographical regions show that differential invasiveness of Acacia species is not always determined by a broad promiscuity with rhizobial strains (Klock et al. 2016; Keet et al. 2017). Similar approaches could be taken with *L. burttii* to further understand the contribution of host range to legume adaptiveness.

### Molecular players restricting the compatibility of legume-rhizobium associations

Root nodule symbiosis encompasses different checkpoints where suitable symbionts are scrutinized, from the early infection to development of functional, nitrogen fixing nodules (Walker et al. 2020). The rhizobial host range is determined by the perception of specific root flavonoids along with certain rhizobial effectors and genes that contribute to NF and exopolysaccharide production (Sugawara et al. 2018; Acosta-Jurado et al. 2016a; Acosta-Jurado et al. 2019, 2020; Kusakabe et al. 2020; Ratu et al. 2021). From the plant side, the first step in symbiotic partner discrimination is the recognition of specific NFs by NF receptors (Radutoiu et al. 2003; Amor et al. 2003; Smit et al. 2007). The next level of selectivity is imposed by scrutiny of expolysaccharides (EPS) produced by rhizobia. In Lotus, this relies on the EPS receptor *LjEpr3* (Kawaharada et al. 2015) and EPS signalling appears to be of general importance across legumes. The incompatibility of *S. meliloti* Rm41 with *M. truncatula* A17 is abolished by incorporating the succinoglycan-coding *exo* gene of the compatible *S. meliloti* 1021 (Simsek et al. 2007). Similarly, EPS composition confers different levels of rhizobial resistance towards the antimicrobial *M. truncatula* nodule-specific cysteine-rich peptides (NCRs) produced in the nodule cells of certain legumes to impose terminal differentiation of bacteroids (Montiel et al. 2017; Arnold et al. 2018). In this regard, the presence of a functional NCR allele in the *M. truncatula* A17 accession restricts its symbiotic association with *S. meliloti* Rm41. This incompatibility is not present in the *M. truncatula* DZA315 accession that possesses a non-functional NCR allele (Yang et al. 2017; Wang et al. 2017).

The broad host range in *L. burttii* is not explained by any of the plant regulators mentioned above. Unlike legumes of the inverted repeat-lacking clade, where terminal differentiation of bacteroids is orchestrated by NCRs, this peptide family is absent in *Lotus* spp. (Kereszt et al. 2018). Our QTL analyses with data generated from 18 Gifu x *L. burttii* RILs inoculated with 12 diverse rhizobial strains showed that a locus near microsatellite marker TM0002 confers the symbiotic promiscuity of *L. burttii*. It is unlikely that *LjEpr3* is responsible for the extended nodulation capacity of *L. burttii*, since this gene is not located near TM0002. By contrast, the NF receptor gene *Nfr1* was an obvious candidate, since it is located near the TM0002 locus. However, Gifu plants expressing a functional extracellular domain of *L. burttii Nfr1* did not result in host-range expansion to include nodulation with *S. fredii* HH103. Since *Nfr1* expression was driven by the Gifu *Nfr1* promoter in this experiment and since there may be additional Gifu/*L. burttii* polymorphisms in other *Nfr1* domains, differences in gene sequence or in endogenous *Nfr1* expression levels between *L. burttii* and Gifu may still account for the difference in nodulation phenotype. However, the *L. burttii* symbiotic associations are established with rhizobial strains producing NF with very diverse decorations (**Table 1**) (D’Haeze and Holsters 2002; Bek et al. 2010; Renier et al. 2011), suggesting that minor changes to nod factor receptors may not be the most likely cause of *L. burttii* promiscuity. An alternative explanation is that other components linked to the TM0002 marker are responsible, for instance, the presence of several resistance proteins in *G. max* restrict strain specific interactions with rhizobia (Walker et al. 2020). Likewise in *Lotus* accessions it was recently found that *Bradyrhizobium elkanii* USDA61 mutants disrupted in different effector proteins of the type III secretion system are affected at different checkpoints in their symbiotic association with *L. burttii, L. japonicus* Gifu and *L. japonicus* MG-20 (Kusakabe et al. 2020). However, the broad host ability of *L. burttii* is unlikely to be linked to a missing R-gene, since the F1 progeny of *L. burttii* and Gifu crosses retained the nodulation capacity with *R. leguminosarum* Norway, while Gifu wile type plants were unable to develop nodules. Therefore, the genetic components responsible for the pronounced symbiotic promiscuity of *L. burttii* remain elusive.

## Conclusion

In this study we have shown that *L. burttii* exhibits a remarkably broad host range, which is controlled by a single, dominant genetic locus near the TM0002 marker.

## Author contributions

M.Z., N.S., H.J., Y.-Y.L., E.J. performed experiments. J.S., S.U.A., M.P., M.M. provided resources. M.Z., and J.M. analysed data. S.U.A. supervised the project. M.Z., J.M. and

S.U.A. wrote the paper.

## Acknowledgements

This work was funded by Danish National Research Foundation grant DNRF79.

---

**Supplementary Figure S1.**
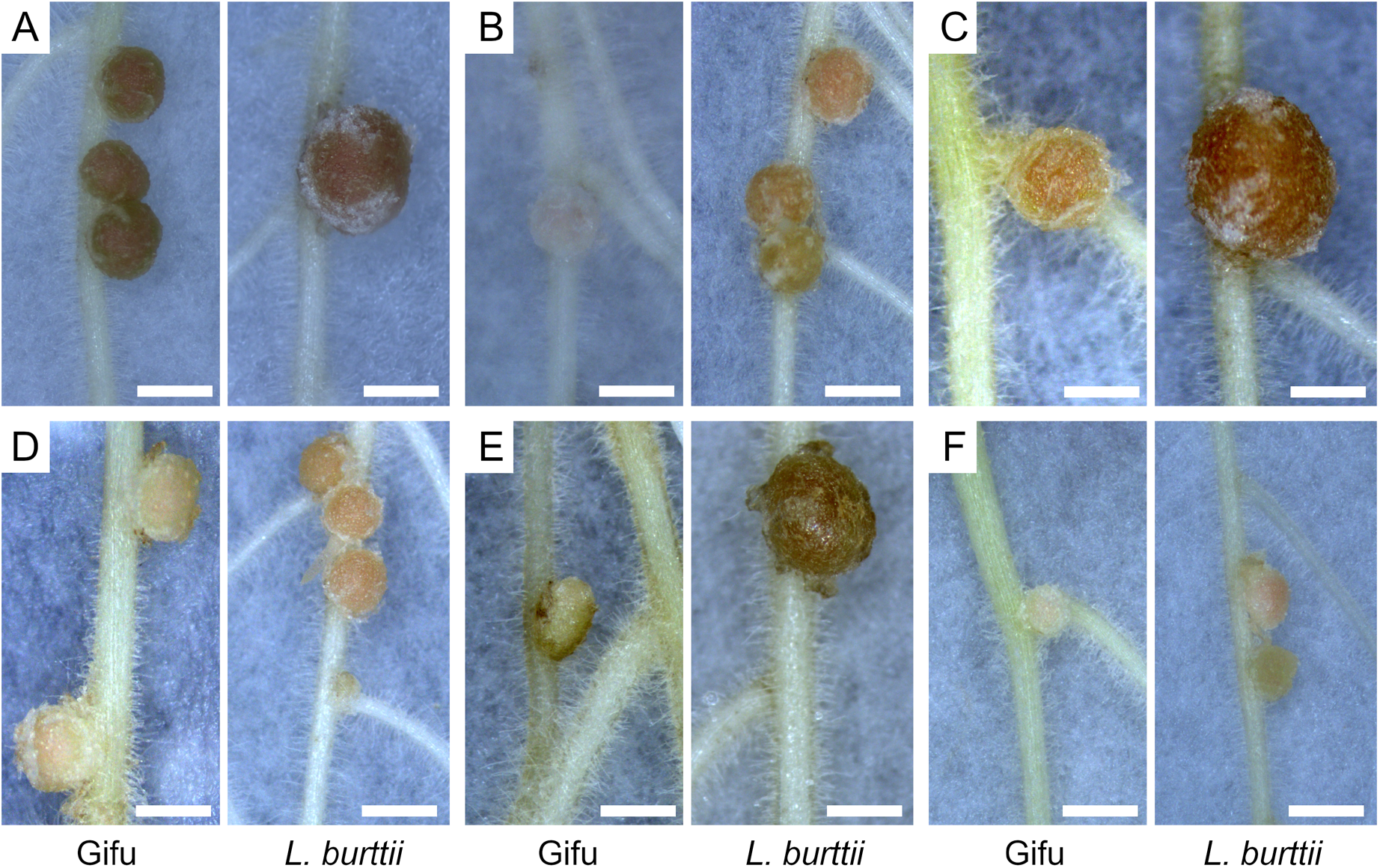
Contrasting nodulation phenotypes in Gifu and *L. burttii* with different rhizobial species. Representative images of nodule structures developed in Gifu and *L. burttii* plants at 5 wpi with *S. fredii* NGR234 (A), *S. terangae* LMG7834 (B), *R. leguminosarum* SM20 (C), *R. leguminosarum* SM5 (D), *M. plurifarium* PMS0804 (E) and *B. pachyrhizi* PMS0802 (F). Scale bar, 1 mm.

